# Increasing age is independently associated with higher free water in non-active MS brain - A multi-compartment analysis using FAST-T2

**DOI:** 10.1101/2021.02.01.429067

**Authors:** Liangdong Zhou, Yi Li, Xiuyuan Wang, Elizabeth Sweeney, Hang Zhang, Emily B. Tanzi, Jennette Prince, Victor Antonio Su-Ortiz, Susan A. Gauthier, Thanh D. Nguyen

## Abstract

**Purpose:** To explore the relationship between the cerebral cortical perivascular space (PVS) and aging in non-active MS subjects by using the multi-echo T2 relaxometry based cerebrospinal fluid fraction (CSFF) map.

**Methods:** Multi-echo spiral T2 data from 111 subjects with non-active multiple sclerosis (MS) were retrospectively investigated by fitting the T2 data into a three-compartment model, the three water compartments including myelin water, intra-extracellular water, and cerebrospinal fluid. Segmentation of T1w image was performed to get the region of interest (ROI) in cerebral cortical regions. The white matter lesion segmentation was conducted using a convolutional neural network (CNN) based segmentation tool. The CSFF in the ROIs were correlated with age by controlling the gender, white matter hyperintensity lesion burden, and MS disease duration. Multiple linear models were created for the analysis of aging effect on the CSFF.

**Results:** The ROI analysis shows that the CSFF in the cerebral cortical regions (temporal, occipital, parietal, front, hippo, and mtl) are significantly linear increasing with age (p<0.01). The intra-extracellular water fraction (IEWF) in the ROIs are significantly linear decreasing (p<0.01).

**Conclusion:** The multi-echo T2 based three-compartment model can be used to quantify the CSFF. The linear increase of CSF water contents in the cerebral cortical regions indicates increased perivascular space load in cortex with aging. The quantification of CSFF may provide a way to understand the glymphatic clearance function in aging and neurodegenerations.

**Highlights:**

- MR T2 relaxometry is a valid method to quantify the cerebrospinal fluid fraction (CSFF) in cerebral cortical regions
- The CSFF in the cerebral cortical regions are positively correlated with age by controlling the white matter lesion load in non-active MS subjects.
- Quantification of cerebral CSFF may reflect the perivascular space load in cortex and better interpret the disease progression in neurodegenerative disease, such as MS.

## 1. Introduction

Dilated perivascular space (PVS) (1), or Virchow-Robin (VR) spaces, is common radiology finding in clinical practice and brain research. PVS surround the walls of vessels as they penetrate through the brain parenchyma from the subarachnoid space, which has been verified functioning as brain clearance pathway of the glymphatic system in recent studies (2–4). However, only enlarged PVS is visible in clinical MRI examination, and those along small vessels in cerebral cortex are hard to see in regular MRI sequence (5). There are studies on the scoring system for qualitative evaluating of the PVS (5). The qualitative scoring system categorize the subjects into 5 groups by counting the PVS number on the slice of semioval. This scoring system could be inaccurate due to the poor image quality, and the unequal distribution of the PVS across image slice. There are also quantitative method for evaluating the PVS volume (6–8). They are doing the imaging processing, such as image enhancement, segmentation, based on the multimodal MRI. Therefore, the performance of the methods also highly relies on the image quality. Most importantly, imaging processing-based methods have an inherent disadvantage for PVS evaluating, that is they are not able to handle the invisible PVS that are not shown in the image, no matter qualitative or quantitative methods. It has been shown that the CSF water accumulates in the PVS (9,10). This provides us an alternative way to measure the PVS by quantifying the CSF water. T2 relaxometry based three water compartment model has been applied in brain research for quantitative water fraction measure, but all of them focused on the myelin water in multiple sclerosis (MS) study (11–13).

MS is a complex disease, characterized by inflammatory demyelination and axonal degeneration. It is hard to treat because the etiology is unknown. A better understanding of the disease mechanism will facilitate the development of new therapies (14). Aging is the most common cause of neurodegenerations. We utilized aging as a risk factor of neurodegenerations and tested its relationship with brain parenchyma CSF fraction (CSFF) in non-active MS brain. CSFF has been hypothesized as a potential biomarker of perivascular space (PVS) of vessels, which has been verified functioning as brain clearance pathway of the glymphatic system in recent studies (2–4). We hypothesize that cerebral cortex CSFF will increase with aging in non-active MS independent of white matter lesion load (neuroinflammation). T2 relaxometry-based multiple water-compartment method has been applied in brain research for quantitative water fraction measure 15), but most of them focused on the myelin water in multiple sclerosis study (11–13). In this study, we first applied the spiral trajectory T2 relaxometry method described in (15) and the three-pool water compartment model (13) in cerebral cortical regions to quantify the CSF water fraction and investigated the aging effect on PVS. We hypothesize that this method will improve the capability of detecting the PVS change in aging and neurodegeneration disease.

## 2. Material and methods

### 2.1 Theory

The three water compartment model has been used in the myelin water quantification in the MS study (13,16,17). In this study we follow the similar procedure as in (13) for the quantification of CSF water. According to the T2 relaxation time difference, the brain water can be modeled as three compartments: the myelin water with T2 time about 10 ms, the intra-extracellular water with T2 time about 50-80 ms, and the CSF water or free water with T2 time larger than 1000 ms at 3T (18). Therefore, the measured MR signal *S*(*t*) in three-pool water compartment model can be written as

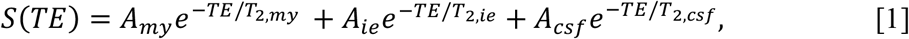

where *TE* is the echo time, *A*_*my*_, *A*_*ie*_, and *A*_*csf*,_ are the signal amplitude attribute to the three water compartments, *T*_2,*my*_, *T*_2,*ie*_ and *T*_2,*csf*_ are the T2 time of the three water compartments, respectively.

The measured signal can then be fitted to the model [1] using nonlinear least square (16) with multiple *TE* s to obtain the solution ***x*** *=* (*A*_*my*_, *A*_*ie*_, *A*_*csf*_, *T*_2,*my*_, *T*_2,*ie*_, *T*_2,*csf*_):

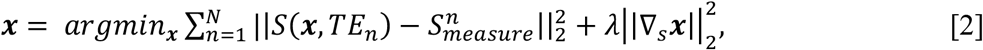

where *S*(***x***, *TE* _*n*_) is the signal model in [1] with *n*-th TE, *N* is the total number of TE, 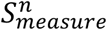 is measured signal at *TE* _*n*_, *λ* is the regularization parameter to impose a spatially local smoothness of the solution (19), and *∇*_*s*_ is the 2D discrete Laplace operator. The regularization parameter was selected by testing on the healthy subjects with various *λ* and chose the one that generated myelin water map with the best visual quality.

Then the CSF water fraction (CSFF) map can be calculated as:

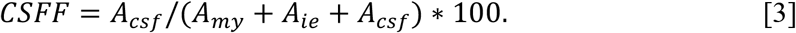

### 2.2 Subjects

This was a retrospective study. 111 non-active MS patients (male: 33, female: 78, age: 58.02±9.70 years, maximum age: 79, minimum age: 45) were scanned as part of a large imaging research database for MS disease. The 111 subjects were filtered by using the following three conditions to avoid the effect of progression of the disease on our purpose:

1. MS is non-active,
2. Have diagnosis, MRI and clinic visits,
3. The gap of diagnosis and MRI visits is less than 12 months.

The details of the filtered subjects’ information are listed in the Table 1.

**Table 1.** Basic information of the subjects.

|  | Female | Male | Total |
| --- | --- | --- | --- |
| **No. of subject** | 78 | 33 | 111 |
| **Age (SD)** | 57.42 (9.47) | 59.42 (10.22) | 58.02 (9.70) |
| **DD (SD)** | 17.56 (9.46) | 17.91 (9.29) | 17.66 (9.37) |
| **MSLB (SD)** | 13.04 (15.04) | 10.64 (15.42) | 12.33 (15.12) |

### 2.3 Data acquisition

Multi-echo T2 data was acquired with Fast Acquisition with Spiral Trajectory and adiabatic T2prep (FAST-T2) sequence at 3T. The FAST-T2 imaging parameters were as follows: axial field of view = 24 cm; matrix size = 192 × 192 (interpolated to 256 × 256); slice thickness = 5 mm; number of slices = 32; spiral TR = 7.8 ms; spiral TE = 0.5 ms; number of spiral leaves per stack = 32; flip angle = 10°; readout bandwidth = ±125 kHz; TEs = 0, 7.6, 17.6, 67.6, 147.6, 307.6 ms. Corresponding T1w, T2w, and T2FLAIR were also acquired at the same session for the anatomical structure and disease diagnosis.

### 2.4 Data processing

CSFF map was computed from multi-echo T2 data acquired with Fast Acquisition with Spiral Trajectory and adiabatic T2prep (FAST-T2) sequence at 3T. A three-compartment nonlinear least squares fitting algorithm with spatial regularization was used to derive the water fraction maps, with the component with the longest T2 assigned to CSF.

FreeSurfer was used on T1w for the segmentation of brain ROIs: frontal lobe, temporal lobe, occipital lobe, parietal lobe, hippo, and medial temporal lobe. The ROIs then were coregistered to the CSFF space and visually checked by three experienced radiologists for the accuracy of coregistration. CSFF in the ROIs was extracted for all subjects.

MS lesion was segmented using in-house developed model based on convolutional neural network with information of T1w, T2w, and T2FLAIR images. The absolute MS lesion volume was extracted from the segmentation. The skull size scaling factor was processed using FSL toolbox. The MS lesion burden (MSLB) was then computed by multiplying absolute volume with the skull size scaling factor for each subject. The skull size scaling factor was used for the normalization of different head size of subjects.

Eighty subjects without obvious MS lesion in the brain centrum semiovale region (BCS) were selected for the PVS score test. Three experienced neuroimaging researchers rated the PVS score of BCS in T2w image individually and arrived at a consensus PVS rating score for each subject by using a published method (5): 0 (none), 1 (1-10), 2 (11-20), 3 (21-40), 4 (>40).. The CSFF in the white matter of the same region was extracted.

The disease duration (DD) is defined as the time gap between MRI date and diagnosis date.

### 2.5 Data analysis

As our goal is to analyze the relationship between CSFF and age, we have to control the effect of other factors such as the gender, DD, and MSLB. We use multivariates linear regression model for the data analysis. CSFF is the output and age, gender, DD, MSLB are the inputs of the model. Variance inflation factor (VIF) was computed for all variables to make sure there is no collinearity between variables. It turned out that the VIF for all variables are close to 1, which indicates there is no collinearity between variables. The cross correlation of variables was calculated to check the interactive effects. The distribution of MSLB is nonGaussian so that we have used a log transform to obtain a nearly normal distributed MSLB. The following descript the model for the two considered outputs:

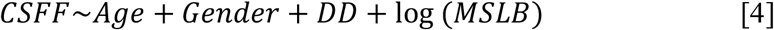

IEWF was not analyzed separately because CSFF+IEWF+MWF=1 and MWF is very small compare with CSFF and IEWF. The trends of IEWF with age is opposite to CSFF.

For the relationship between WM CSFF and PVS score on the semiovale slice, we fitted an ANOVA model.

To see how the DD affects the CSFF within the non-active MS cohort, we filtered our subjects with two conditions: one is subjects age>=50 years old, the other one is grouping the filtered subjects using the first condition into two groups (DD_long and DD_short) by DD. The cutoff DD between the two groups is the median of DD (15 years), 41 subjects for each group.

### 2.6 Code and data availability

The code and data are available with a request sending to the corresponding author by following the data sharing restrictions in our institute.

## 3. Results

### 3.1 CSFF with PVS score

We fit an ANOVA model and found that there was a difference in WM CSFF by PVS score (F = 5.83, p-value < 0.001), (Figure 1). This implies that CSFF can serve as an alternative indicator of PVS change. Interestingly, CFFS doesn’t seem very different for PVS score 0-2, but starts to increase at a score of 3. The higher data variability of CSFF in groups 1 and 2 were observed.

**Figure 1.**
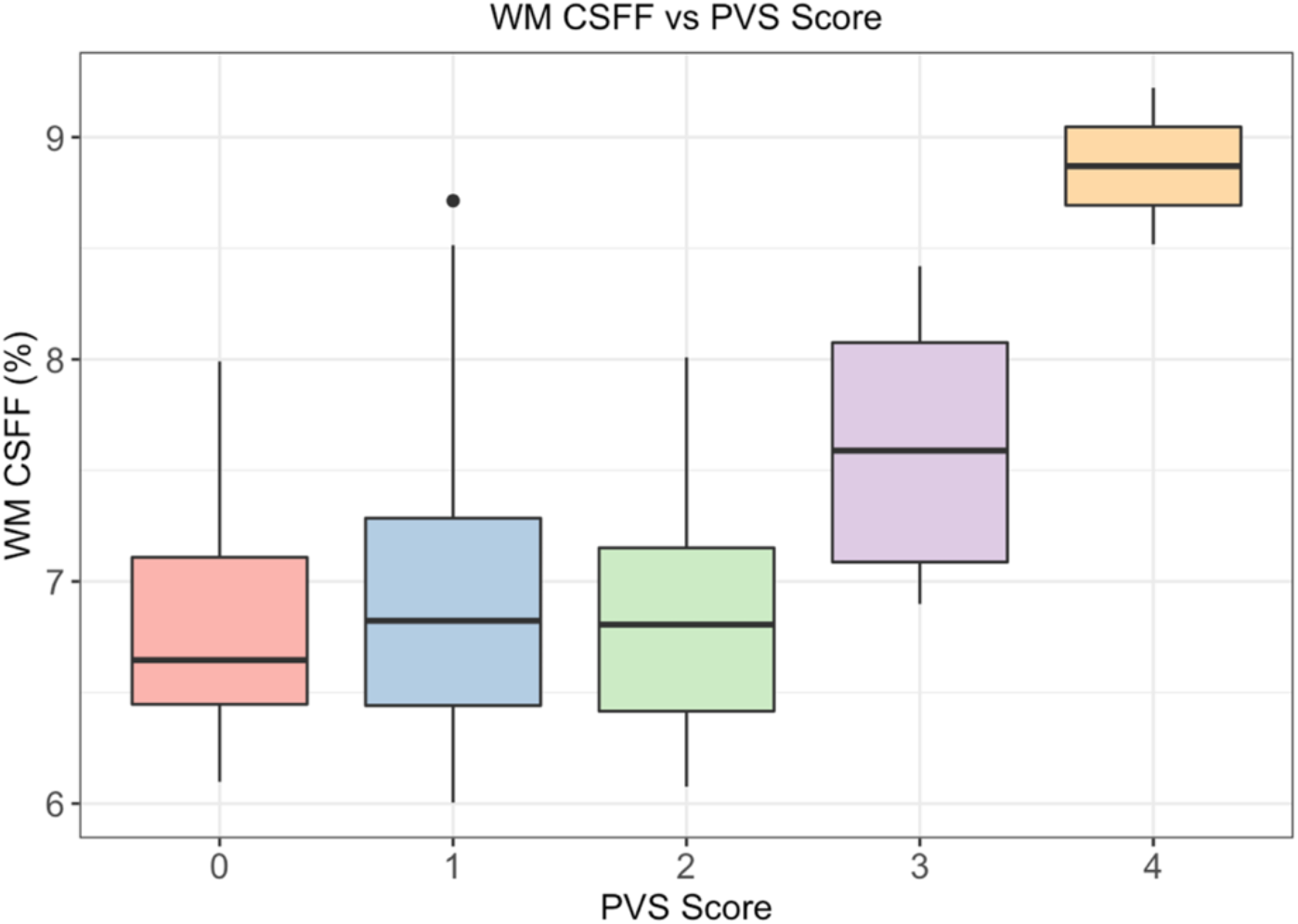
WM CSFF and PVS score of centrum semiovale white matter.

### 3.2 CSFF with MSLB in WM

According to multiple simple linear regression analysis, it shows in Figure 2 that CSFF in all selected brain regions are significantly (*p <* 0.5) linear with the total white matter lesion burden.

**Figure 2.**
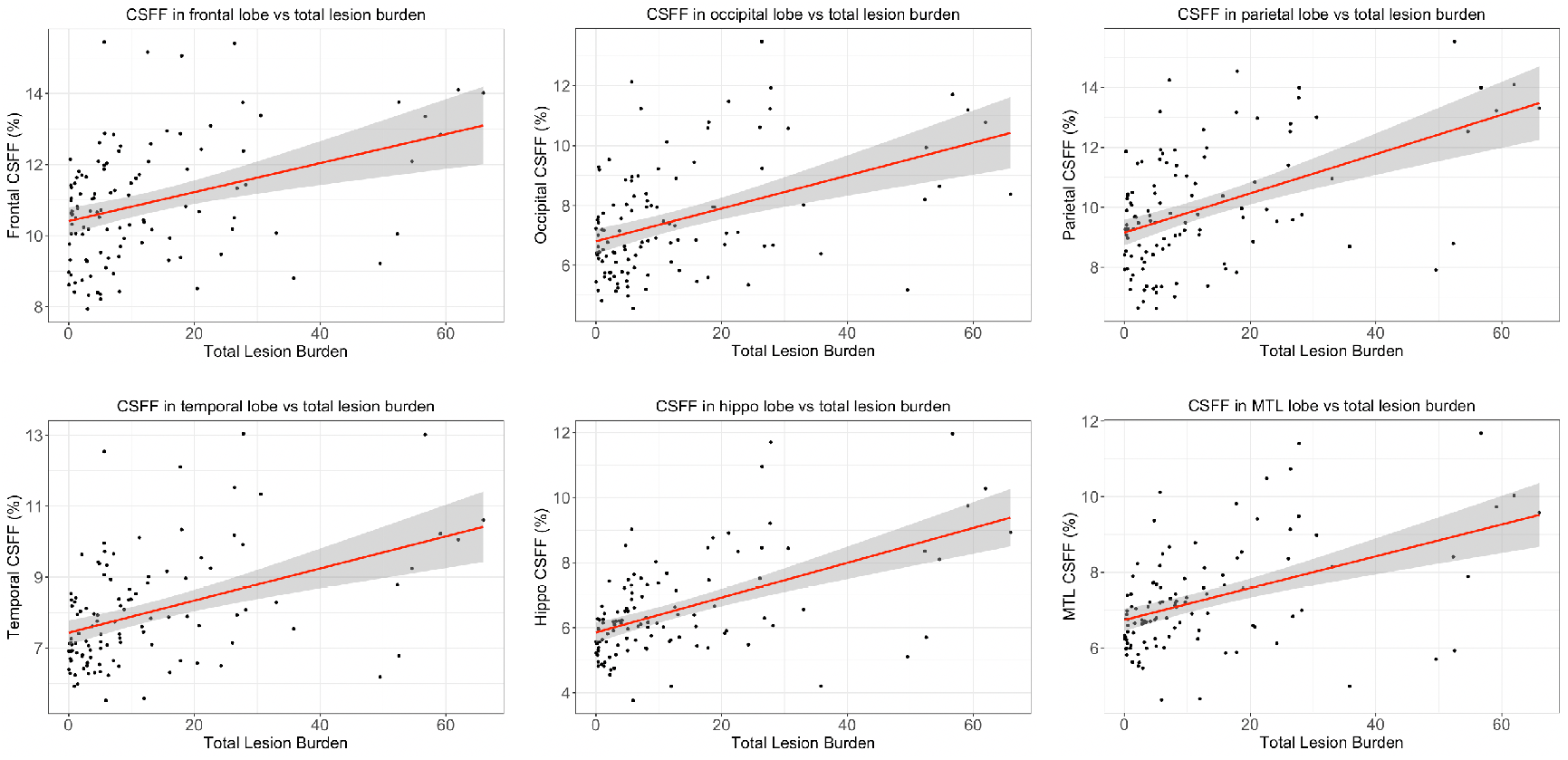
CSFF versus white matter lesion burden in selected brain regions. It shows that CSFF is linear with WM lesion burden in all selected regions with p<0.01.

### 3.3 White matter lesion with age

The linear relationship between the total white matter lesion burden and age is shown in Figure 3. This results is previously observed in literature (20,21).

**Figure 3.**
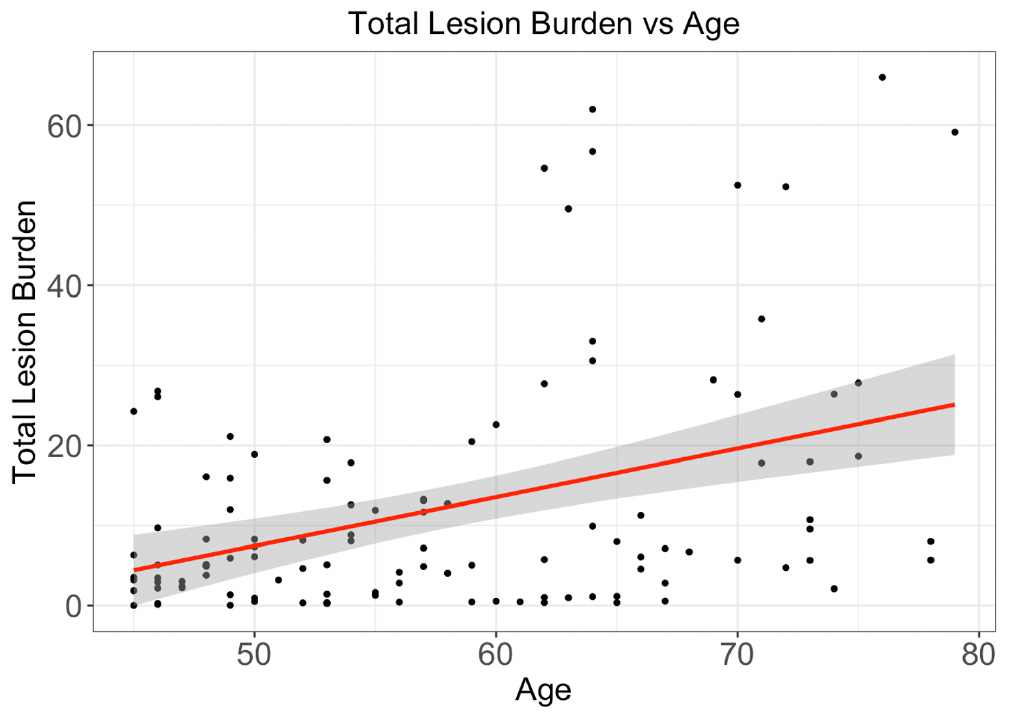
White matter lesion burden with age. It shows that the WM lesion burden is linearly correlated with age (p<0.01).

### 3.4 CSFF with disease duration

The relations between CSFF and disease duration is presented in Figure 4. It shows the significant linear relationship between these two values.

**Figure 4.**
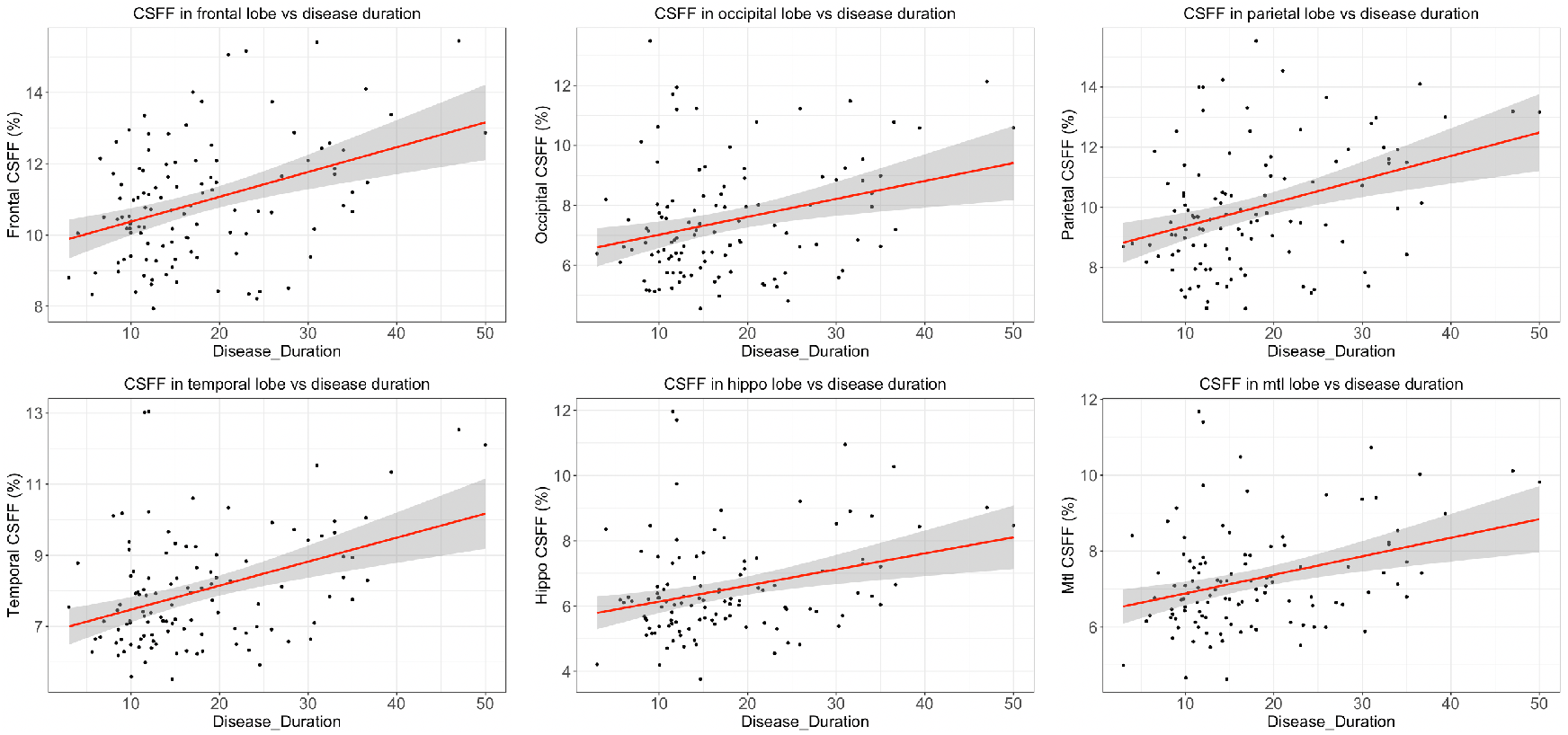
CSFF with disease duration. It shows that CSFF is significantly linear with the disease duration in all the selected brain regions (p<0.01).

### 3.5 CSFF with multiple variates analysis

The VIF for variables are presented in Table 2. It shows that the VIF for all considered variables are close to 1, which is an indicator of absence of collinearity. Therefore, we can safely build a linear model to investigate the factors that have an effect on the output variables, CSFF, IEWF.

**Table 2.** VIF table for variables to determine if there is a collinearity between variables.

| VIF table for variables |  |  |  |  |
| --- | --- | --- | --- | --- |
| Variables | Age | Gender | DD | MSLB |
| VIF | 1.2606 | 1.0288 | 1.1317 | 1.1656 |

The results in Figure 5 show that cerebral cortical CSFF significantly correlated with age in all brain lobes after controlling for gender, and for disease factors such as DD and MSLB. Table 3 shows the linear regression coefficients. The p-values in the last column indicates that all the variables have significant contribution to the output CSFF in the temporal lobe. We need to control the variables to extract the pure effects of variable on the output, i.e., we need to do the p-value adjustment by controlling the variables. The p-value adjustment was done by utilizing the false discovery rate (FDR). The FDR adjusted p-value for temporal CSFF regression model was shown in Table 4. The regression model coefficients and the FDR adjusted p-value table for CSFF in other ROIs similar to Table 3 and Table 4 can be found in the supplementary document.

**Figure 5.**
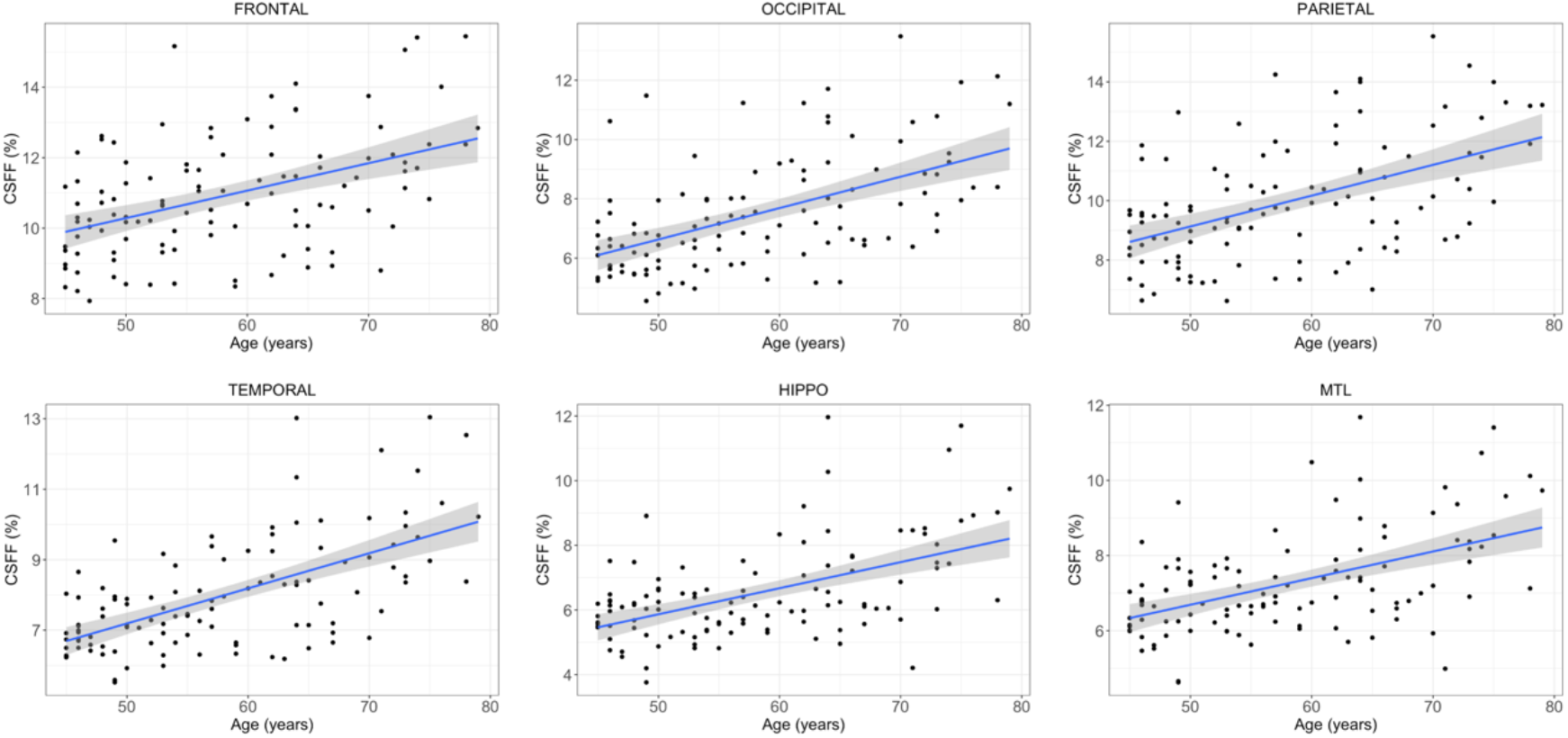
Linear relationship between CSFF and Age in the six regions.

**Table 3.** Multivariable analysis of csff in temporal lobe with Age, gender, DD, and mslb.

|  | Estimate | Std. Error | t value | p value |
| --- | --- | --- | --- | --- |
| **(Intercept)** | 2.3286 | 0.6513 | 3.576 | 5.28e-4 *** |
| **Age** | 0.0699 | 0.0124 | 5.625 | 1.52e-7 *** |
| **GenderM** | 0.635 | 0.2370 | 2.679 | 0.0086 ** |
| **DD** | 0.0338 | 0.0122 | 2.775 | 0.0065 ** |
| **log(MSLB)** | 0.4074 | 0.1046 | 3.894 | 1.73e-4 *** |

Significant codes: 0:\*\*\* 0.001:\*\* 0.01:\* 0.05:.

**Table 4.**
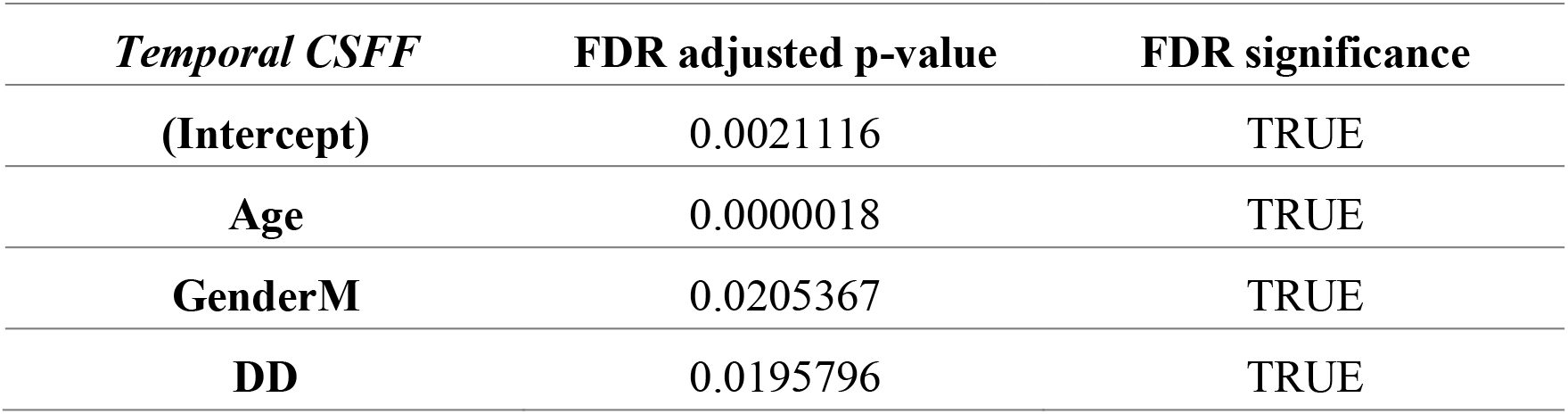
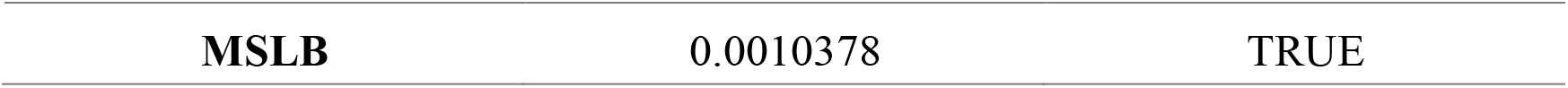
FDR adjusted p-value for temporal CSFF regression model.

### 3.6 CSFF for subjects with long disease duration

Figure 6 presents the boxplots of the two group subjects in CSFF, age, DD, and MSLB. It shows that there is no significant age and MSLB difference between two groups. However, we observe significant difference of CSFF between two groups in the global gray matter, frontal and parietal lobes. And CSFF is marginally different in white matter for two groups.

**Figure 6.**
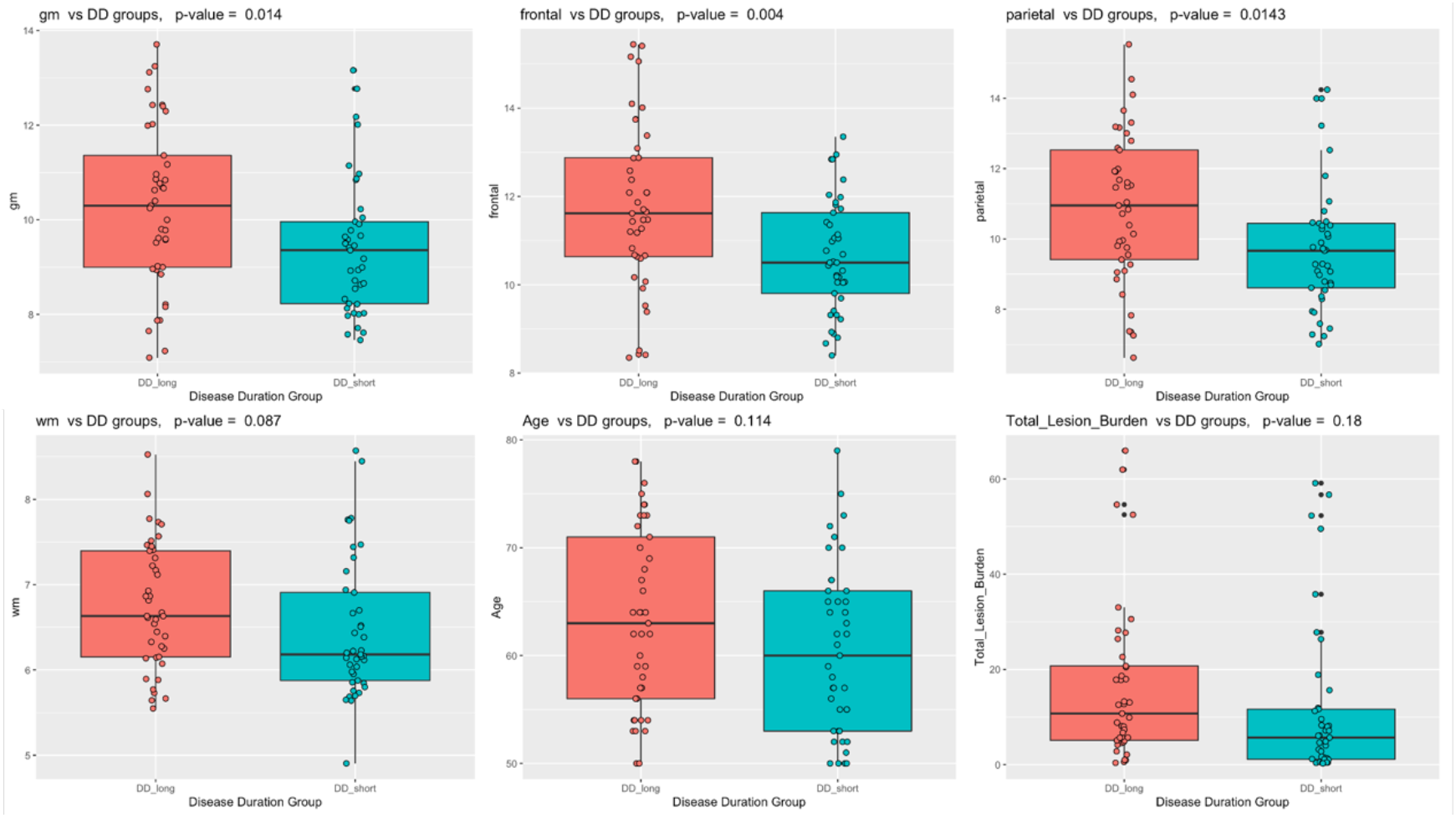
Boxplot of the CSFF, age, DD and MSLB of two groups. It shows that CSFF for two groups are significantly different in globe gray matter, frontal lobe, and parietal lobe, but marginally significantly different in global white matter.

## 4. Discussion

The role of neurodegeneration in MS has got more attention in recent years, which may involve the disfunction of the recent discovered brain glymphatic system (22). PVS is an important pathway of the glymphatic system and has been reported link with aging and neurodegeneration disease. On the other hand, CSFF reflects the nature of CSF water compartment in brain parenchyma and has been hypothesized as a quantitative measure of PVS which was filled with CSF (9,10). T2 relaxometry based modeling provides us an alternative way to measure PVS by evaluating the CSFF. With this method, we observed CSFF increases with aging in cerebral cortex for the first time by controlling MS disease factors, which indicates that CSFF could be a potential biomarker to evaluate PVS function in MS, which is an important pathway of glymphatic clearance.

MRI visible PVSs have been reported to increase with age in both WM and basal ganglia (23,24). This phenomenon has been long suspected in the cerebral cortex (25,26), but no human study has been reported because of a lack of a valid imaging tool. The quantification of CSFF not only includes the visible PVS shown in conventional MR image but also the invisible PVS in conventional MR image based on nature of the MR T2 relaxometry method.

We observed a linear increase of CSFF in cerebral cortex with age after controlling for disease duration and total white matter lesion burden. The diameter of large PVS (>3mm) increasing with age has been observed in the basal ganglia and white matter. We for the first time applied the multi-echo spiral T2 to map the CSF water fraction in cerebral cortex and observed CSF water compartment increased with aging in cerebral cortex in stable MS subjects. The proposed CSFF mapping method solves the problem of invisible PVS quantification in conventional PVS score system and potentially serves as a more accurate PVS quantification method. Glymphatic clearance play a important role is neurodegeneration disease such as AD, However, we need better understand it’s role in MS.

In Figure 1, the PVS score increasing with age has been observed in the brain centrum semiovale region, which agrees with previous report. The CSFF doesn’t seem very different for PVS score 0-2 but starts to increase at a score of 3. This may be because the portion of the visible enlarged PVS is a relatively small part of total PVS in those subjects, and the CSFF measure reflecting total PVS was driven by those invisible small ones. The higher data variability of CSFF in groups 1 and 2 also indicates that varying levels of PVS were included in those groups.

The proposed CSFF quantification method has several advantages and potential applications. First, it is a biophysical based modeling fitting. This is the essential difference between our proposed method and the conventional image based postprocessing. Therefore, the CSFF method is not only able to quantify the free water in the visible PVS from MRI, but also capable of quantifying the free water in the PVS that surround the small vessels and invisible in the MRI. Second, to the best of our knowledge, this is the first study that quantify the CSFF in the cerebral cortex regions. Because the PVS in the cortex region usually presents as tiny or invisible dots on conventional MRI, this approach offers a new way to investigate the PVS load in brain parenchyma.

Moreover, the accurate PVS quantification from CSFF is promising to be applied in the neurodegenerative disease. The CSF is found playing the key role in the waste clearance procedure. An accurate CSF quantification in brain parenchyma might open a door for the mystery of many diseases, such as MS, AD, PD, SVD. For instance, in this study, we found that in elderly subgroup of the non-active MS subjects, the CSFF increase significantly in the frontal and parietal lobes for those subjects with long disease duration but without significant age difference between two groups as shown in Figure 6. This could imply that MS disease of long duration keep affecting the brain in a subtle way, which could be monitored by the quantification of CSFF.

Besides the advantages mentioned above, there are several limitations of this study. First, the data is limited. More data with various type of MS disease could be helpful to explore the behavior of CSFF in a broad perspective. Second, the CSFF could be a biomarker of PVS. However, this hypothesis is not fully validated. We scored the PVS number by visual check of T2w image, which could be biased by image resolution and radiologist experiences. Theoretically, CSFF represents the free water fraction which includes free water in visible PVS and invisible PVS in T2w image. It is our future work to validate the relation between CSFF and PVS using postmortem study.

## 5. Conclusions

We proposed a T2 relaxometry based CSFF quantification method, which is able to quantify the CSF water in brain parenchyma. Analysis shows that the CSFF correlates with PVS score. The linear increase of CSF water contents in the cerebral cortical regions indicates the increased perivascular space load in cortex with aging. The quantification of CSFF could potentially provide a way to understand the PVS function in brain aging and neurodegenerations, which is part of the glymphatic clearance system.

## Acknowledgement

Supported by NIH grants: AG057848.

